# Altered Brain Dynamics in idiopathic REM sleep behavior disorder: Implications for a continuum from prodromal to overt alpha-synucleinopathies

**DOI:** 10.1101/2023.05.05.539548

**Authors:** M Roascio, SH Wang, V Myrov, F Siebenhühner, R. Tro, P. Mattioli, F. Famà, S. Morbelli, M. Pardini, JM Palva, D Arnaldi, G. Arnulfo

**Author notes:** Correspondence to: Monica Roascio, Full address: via all’Opera Pia 13 (16145), Genova (GE), Italy. These authors equally contributed to this work (D Arnaldi, G. Arnulfo).

## Abstract

Idiopathic/isolated REM sleep behavior disorder (iRBD) is considered a prodromal stage of alpha-synucleinopathies. Cortical and sub-cortical brain modifications begin years before the emergence of overt neurodegenerative symptoms. To better understand the pathophysiological process impacting the brain from the prodromal to the overt stage of alpha-synucleinopathy, it is essential to assess iRBD patients over time.

Recent evidence suggests that the human brain operates at an operating point near a critical phase transition between subcritical and supercritical phases in the system’s state space to maintain cognitive and physiological performance. In contrast, a deviation from the critical regime leading to altered oscillatory dynamics has been observed in several pathologies. Here, we investigated if the alpha-synucleinopathy produces a deviation of the operating point already evident in the prodromal phase and if this shift correlates with biological and clinical disease severity.

We analyzed a dataset of 59 patients with iRBD (age 69.61 ± 6.98, 50 male) undergoing resting-state high-density EEG, presynaptic dopaminergic imaging, and clinical evaluations. Thirty-one patients (age 72.41 ± 7.05, 31 male) also underwent clinical and instrumental follow-up (mean follow-up period 25.85 ± 10.20 months). To localize the individual operating points along the excitation-inhibition (EI) continuum, we assessed both measures of neuronal EI balance and measures of critical brain dynamics such as long-range temporal correlation (LRTCs) and neuronal bistability in spontaneous narrow-band oscillations. Finally, we correlated critical brain dynamics and EI balance metrics with phase synchronization, nigro-striatal dopaminergic functioning, and clinical performances.

Compared to 48 healthy subjects (age 70.25 ± 10.15, 23 male), iRBD patients showed higher values of LRTCs and bistability in the 2-7 Hz band at diagnosis. Patients who eventually phenoconverted to overt alpha-synucleinopathy exhibited a more excitation-dominated (fEI > 1) condition than stable iRBD patients in 5-7 Hz. This higher excitation also directly correlated with phase synchronization in 2-7 Hz, further suggesting a shift of the operating point toward a supercritical state with the disease progression. Moreover, excitation-dominated state and low bistability were associated with deterioration of the nigro-striatal dopaminergic function and tended to correlate with stronger clinical symptoms.

In conclusion, this study shows for the first time a deviation of the working point from inhibition-to excitation-dominated states along the continuum from prodromal to overt phases of the disease. These cortical brain dynamics modifications are associated with nigro-striatal dopaminergic impairment. These results increase our knowledge of the physiopathological process underlying alpha-synucleinopathies since prodromal stages, possibly providing new clues on disease-modifying strategies.

## Introduction

Alpha-synucleinopathies are neurodegenerative diseases involving cortical and subcortical structures, characterized by highly disabling symptoms. The main clinical entities are represented by Parkinson’s disease (PD), dementia with Lewy bodies (DLB), and multiple system atrophy (MSA). Clinical diagnosis generally occurs in the presence of overt syndromes including parkinsonism and/or dementia. However, the neurodegeneration process begins several years prior to the emergence of overt neurological symptoms. For example, when PD is diagnosed about 50% of dopaminergic neurons are supposed to be already lost (Shapira et al., 2017). Most literature has focused on brain modifications in the overt phase of alpha-synucleinopathies, while longitudinal data on the prodromal phase (before the diagnosis of the overt phenotypes) are limited. Exploring the longitudinal changes of cortical and subcortical structures since the prodromal phase of alpha-synucleinopathies can increase our knowledge of the neurodegeneration process since the earlier stages, possibly providing clues for new disease-modifying treatment strategies. REM sleep behavior disorder (RBD) is considered the strongest biomarker of the alpha-synucleinopathy in prodromal phase (Heinzel et al., 2019; Postuma et al., 2019). Thus, patients with idiopathic/isolated RBD (iRBD) represent an ideal population to longitudinally evaluate the brain modifications occurring before the emergence of the full-blown neurodegenerative disease.

The human brain operates near the boundary of a critical phase transition between subcritical and supercritical phases in the system state space (Chialvo, 2010; O’Byrne & Jerbi, 2022; Zimmern, 2020). Shifts of the operating point along the excitation/inhibition continuum track changes in the emerging internal dynamics and are tightly linked with the system working conditions. Supercritical or subcritical states indicate excessively excitable or inhibited pathological conditions such as epilepsy or coma, respectively (Cerf et al., 2004; Liu et al., 2014). However, contrary to simpler dynamical systems (*e*.*g*., interacting water molecules), recent modeling and empirical evidence demonstrate that the phase transitions of the human brain can be both continuous and discontinuous, showing a high degree of bistability (Wang et al., 2022).

Synchronization of neuronal oscillations is crucial for information transformation and for optimizing information processing, transmission, and storage (Fries, 2005, 2015; Hahn et al., 2019). Hence, synchronous oscillations represent a possible controlling mechanism to regulate shifts in the working regime. Indeed, deviation from the optimal critical regime is associated with alterations in physiological oscillatory dynamics in different brain disorders, such as epilepsy (Fusca et al., 2022; Linkenkaer-Hansen et al., 2005; Wang et al., 2022), autism spectrum disorder (Bruining et al., 2020), and neurodegenerative diseases (Montez et al., 2009). However, few studies have investigated whether these dynamics are already altered in the prodromal stages of neurodegenerative disease (Javed et al., 2022; Maestú et al., 2021).

Previous studies of iRBD have mainly investigated how large-scale couplings of brain oscillation change over time (Roascio et al., 2022) or differ between patients and healthy controls (Sunwoo et al., 2017). Patients with iRBD exhibit weaker synchronization compared to age-matched healthy subjects in the delta band (Sunwoo et al., 2017) at diagnosis. Phase synchronization further increases in the alpha band following disease progression (Roascio et al., 2022). However, observing changes in spontaneous oscillatory dynamics alone does not inform the direction of the operating point’s shift. Combining them with measures of excitation-inhibition (EI) balance, bistability, and long-range temporal correlations (LRTCs) could provide direct evidence of the direction of shift of the operating point.

In the present study, we hypothesized that iRBD patients show altered critical brain dynamics with the operating point shifted to excitation-dominated states, similar to how mild cognitive impairment (MCI) affects brain dynamics before Alzheimer’s dementia (Javed et al., 2022; Maestú et al., 2021; Pusil et al., 2019). We also expected the extent of these alterations to be highly correlated with disease severity, as assessed by both instrumental and clinical biomarkers. To this end, we first investigated how LRCTs, EI balance, and brain bistability changed in iRBD patients compared to healthy controls in resting-state high-density EEG data. Then, we observed how the critical dynamics change in iRBD patients who remained idiopathic with disease progression, compared to iRBD patients who will phenoconvert/phenoconverted into alpha-synucleinopathies over time. Secondly, we studied how critical brain dynamics correlated with the biological and clinical severity of neurodegeneration in iRBD patients. To this aim, we correlated the features extracted from the EEG time series with (i) nigro-striatal dopaminergic function, as evaluated by [^123^I]FP-CIT-SPECT, and (ii) with clinical scores of motor and cognitive functions. Finally, we investigated whether EI balance is positively correlated with phase synchronization in iRBD patients.

## Materials and Methods

### Sample characteristics

The study was conducted according to the declaration of Helsinki, and all participants gave informed consent before entering the study, which the local ethics committee approved.

The diagnosis of iRBD was confirmed by overnight videopolysomnography, according to current criteria (Sateia, 2014). All patients underwent baseline high-density 64-channels EEG, [^123^I]FP-CIT-SPECT, and a comprehensive clinical assessment including (1) the Mini-Mental State Examination (MMSE) as a global measure of cognitive impairment; (2) the Movement Disorder Society-sponsored revision of the Unified Parkinson’s Disease Rating Scale, motor section (MDS-UPDRS-III) to evaluate the presence of parkinsonian signs; (3) clinical interviews and questionnaires for activities of daily living (ADL) and instrumental ADL to exclude dementia.

All iRBD patients underwent general and neurological examinations to exclude other neurological and psychiatric disorders. Brain magnetic resonance imaging (MRI), or computed tomography in the case MRI was unfeasible, was used to rule out main non-degenerative brain diseases such as tumors or vascular lesions. White matter hyperintensities were not an exclusion criterion if the Wahlund scale was not >1 for each brain region (Wahlund et al., 2001).

All subjects diagnosed with iRBD underwent high-density EEG (hdEEG) evaluation during relaxed wakefulness and brain [^123^I]FP-CIT-SPECT within three months from diagnosis. All patients underwent clinical follow-ups every six months, including motor and cognitive assessments. The occurrence of phenoconversion of PD, DLB, or MSA was defined based on current clinical criteria (Gilman et al., 2008; McKeith et al., 2017; Postuma et al., 2015).

A subgroup of patients also underwent instrumental follow-up including high-density EEG and [^123^I]FP-CIT-SPECT.

Here, we grouped patients as follows: (i) stable iRBD (sRBD) if patients had not phenoconverted at the last available follow-up; (ii) progressive iRBD (pRBD) if patients were still free from parkinsonism/dementia at the time of instrumental (*i*.*e*., hdEEG and [^123^I]FP-CIT-SPECT) follow-up, but phenoconverted in the near future at subsequent clinical follow-up; (iii) converted iRBD (cRBD) if patients already phenoconverted into an overt alpha-synucleinopathy at the time of instrumental follow-up. In this way, we could study clinical and instrumental characteristics of iRBD patients at initial diagnosis and at three follow-up conditions (stable iRBD, progressive iRBD, and phenoconverted patients), describing four steps along the continuum from prodromal to overt stages of synucleinopathies.

As a control dataset, we included healthy control (HC) subjects who underwent baseline clinical evaluation (*i*.*e*., MMSE) and 64-channel hdEEG recording as part of a previous voluntary program in our institution.

### EEG collection and pre-processing

The EEG recording was carried out late in the morning to minimize drowsiness. We used the Galileo system (EBNeuro, Florence, IT) to acquire bandpass (0.3–100 Hz) signals from 64 electrodes at a sampling rate of 512 Hz. We placed the electrodes according to the 10–10 International System, where the reference electrode and ground were Fpz and Oz, respectively. Electrode impedances were monitored and kept below 5 kOhm. An EEG technician monitored the recording session to maintain a constant vigilance level over the patient, prevent sleep, and preserve a high signal quality across the recording session.

We filtered the time series with a notch filter (order 2) to remove power line noise (50Hz). We rejected the channels with a high percentage of artifacts (number of rejected channels 2.57 ± 2.43, mean ± SD). We used independent component analysis (ICA) and visual inspection to reduce the number of physiological and instrumentation artifacts, such as blinks, lateral eye movements, muscle artifacts, drowsiness, and electrode pop. We filtered the time series with a Finite Impulse Response (FIR) band-pass filter (1–80 Hz, Kaiser window, order 1858). We interpolated bad channels using spline interpolation (kernel size: 4 cm). Finally, we applied the scalp current density (SCD) for all clean sensors with the spline method (lambda 0.00001, order 4, degree 14).

### EEG metrics

We analyzed the broadband SCD time series with a time-frequency decomposition using 30 log-spaced narrowband Morlet wavelets (m=5) in the 2-70 Hz range (Tallon-Baudry et al., 1998; Torrence & Compo, 1998). For each time-frequency series, we estimated the detrended fluctuation analysis scaling exponent, the functional EI ratio, the bistability index, and the phase synchronization methods described below.

#### Detrended Fluctuation Analysis

To investigate LRCTs, we performed detrended fluctuation analysis (DFA). We quantified the scale-free decay of temporal correlations in the amplitude modulation of neuronal oscillations (Linkenkaer-Hansen et al., 2001; Poil et al., 2012) with the DFA scaling exponent, which is the slope of the fluctuation function. A DFA exponent in the range [0.5, 1] indicates power-law scaling behavior and the presence of LRCTs, while an exponent below 0.5 is characteristic of an uncorrelated signal.

#### Functional Excitation-Inhibition Ratio

To quantify the EI balance, we compute the functional EI ratio (fEI) for windows with fixed length of 40 cycles for each narrow-band signal and 80% overlap (Bruining et al., 2020). Systems operating at the critical point should have fEI = 1, while those working in the inhibition- or excitation-dominated states should have an fEI < 1 or fEI > 1, respectively.

#### Bistability Index

Brain bistability represents the discontinuous transition between asynchronous and totally synchronous activity (Freyer et al., 2011), and high bistability is considered a sign of brain pathology. To evaluate brain bistability, we quantified the bistability index (BiS) of a power time series R^2^ fitting its probability density function (PDF) with a single- and a bi-exponential model. The single-exponent model is defined as

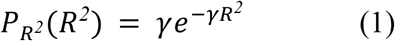

where *γ* is the exponent. The bi-exponential model is defined as

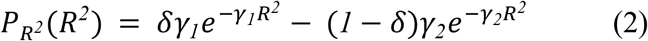

where *γ*_*1*_, *γ*_*2*_ are the two exponents and *δ* is a weighting factor. To assess the fitting of the two models, we used the Bayesian information criterion (BIC)

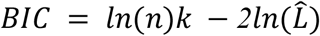

where *n* is the number of samples; 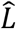 is the likelihood function; *k* is the number of free parameters in the model, i.e., for eq. (1), k = 1 and for eq. (2) and k = 3. A better-fitted model yields a small BIC value.

Next, we computed the difference in BIC between single- and bi-exponential fitting:

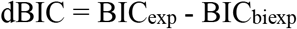

Finally, the BiS is computed as

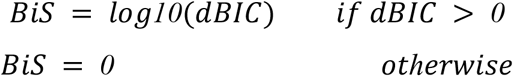

A BiS close to zero means that the single-exponential model is a more likely model for the observed time series, whereas, for a BiS greater than 0, the most likely model for the observed time series is the bi-exponential model.

#### Weighted Phase Lag Index

To quantify the phase synchronization, we computed the weighted phase lag index (wPLI) for each pair of channels (Vinck et al., 2011). The wPLI is neither inflated by volume conduction as the phase-locking value (PLV) nor underestimating the true phase coupling as the imaginary part of the complex-valued PLV (Palva & Palva, 2018) and thus is ideal for EEG sensor level synchrony analysis intended here. The wPLI is limited to the range [0,1], whereas 0 represents the absence of synchronization.

The brain network graph model represents pairwise relations between sets of neuronal elements (*i*.*e*., nodes) that interact throughout pairwise relations (*i*.*e*., links) (Bullmore & Sporns, 2009). These pairwise relations can represent the coherent oscillatory activity between neurons (*i*.*e*., functional connectivity) or the anatomical connection between fibers (*i*.*e*., structural connectivity). The wPLI matrix is a weighted undirected graph, where the nodes are EEG channels (n=60), and the links are the wPLI values. A graph G can be characterized by several graph metrics. Here, we quantified three different node metrics - *i*.*e*., eigenvector centrality, clustering coefficient, and strength - for each channel pair of the wPLI matrix for the following analysis. The eigenvector centrality measures the influence of a node in a graph and is quantified as the number of connections a node has with all other nodes. The strength is the sum of the weights of links connected to the node. The local clustering coefficient measures the degree to which the nodes in a graph tend to cluster and is quantified by the fraction of how many possible triplets (*i*.*e*., three nodes connected together) are realized between the neighbors of a node.

### Molecular imaging evaluation

Subjects with iRBD - within three months since diagnosis - underwent [^123^I]FP-CIT-SPECT to measure the striatal dopamine transporter (DaT) density according to European Association of Nuclear Medicine (EANM) guidelines (Morbelli et al., 2020; Nobili et al., 2010). Details on SPECT data acquisition and analysis can be found in the *Supplementary Materials*.

In brief, reconstructed images were exported in the Analyze file format and processed by Basal Ganglia V2 software (Nobili et al., 2013) to compute specific to non-displaceable binding ratios (SBRs). In particular, background uptake (occipital region) was subtracted from putamen or caudate uptake as follows: (putamen/caudate uptake—background uptake)/background uptake to compute SBR values. For subsequent analyses, we computed the mean SBR values between the right and left hemispheres for the caudate and putamen.

### Statistical Analysis

To investigate the modifications of the acquired metrics over time, the difference between phenoconverted and non-phenoconverted iRBD patents, as well as between iRBD patients and HC subjects, we compared groups using the Wilcoxon rank-sum test across frequencies (*n*=30 Morlet wavelets, *α*=5%).

We computed Spearman’s rank correlation coefficient to investigate the relationship between EEG measures (*i*.*e*., DFA, fEI, and BiS) - averaged across channels - and SBR values separately for the putamen and caudate. For the frequency bands with a significant correlation between EEG metrics and SBR values (*p* < 0.05), we further investigated if there was a correlation (Spearman’s rank correlation coefficient) between abnormal EEG measures and nigro-striatal dopaminergic impairment in single channels.

To assess the diagnostic performance of EEG measures, we looked for a threshold value that best differentiates sRBD and pRBD patients at baseline. To do this, we set a range of possible thresholds between -1 and 1 linearly equispaced. For each value in this range, we computed the f1-score, specificity, and sensitivity (Dhamnetiya et al., 2022). Finally, we chose as the threshold the range value that showed the best performance in discriminating patients with stable and progressive iRBD.

In order to evaluate the correlation between continuous variables (*i*.*e*., EEG features) and ordinal variables (*i*.*e*., MMSE and MDS-UPDRS-III), we used Kendall’s coefficient of rank correlation tau-sub-b (Khamis, 2008).

Finally, we explored the correlation between EI balance and phase synchronization - both averaged across channels - using Spearman’s rank correlation coefficient.

For all previous analyses, we corrected the p-values with the Benjamini–Hochberg (BH) method (*α*=0.05) to reduce the false discovery rate.

### Data availability

The data used in this work can be available upon a reasonable request due to privacy issues of clinical data.

## Results

### Demographic, clinical, and imaging data

We enrolled 62 iRBD subjects (9 female; mean age 79.58 ± 7.21 years) and 48 healthy control subjects (23 female, 70.25±10.26) in collaboration with the Sleep Lab of the Clinical Neurology, University of Genoa, IRCCS Policlinico San Martino (Table 1; Figure 1). Thirty-one people with iRBD (all men, mean age 72.41±7.05) underwent a clinical, EEG, and DaT-SPECT follow-up evaluation after 25.79±12.45 months from the diagnosis. Eighteen iRBD patients (4 female, mean age 73.35±6.05) phenoconverted to Parkinson’s disease (n=8) and dementia with Lewy bodies (n=10) after 23.82±18.13 months.

**Table 1.** Main clinical and demographic data of iRBD patients at baseline and follow-up. For each score, we estimated the statistical difference between baseline and follow-up using the Wilcoxon rank sums test.

|  | Baseline | Follow-up | *p* |
| --- | --- | --- | --- |
| Subjects (#) | 59 | 31 | n.a |
| Age (yy) | $69.61 \pm 6.98$ | $72.41 \pm 7.05$ | 0.14 |
| Sex (female/male) | 9/50 | 0/31 | n.a. |
| Phenoconverted (#) | 0 | 8 (4 PD; 4 DLB) | n.a. |
| MMSE | $27.97 \pm 2.19$ | $27.37 \pm 3.02$ | 0.62 |
| MDS-UPDRS-III | $1.78 \pm 3.22$ | $4.93 \pm 9.49$ | 0.07 |
| SBRs caudate | $6.88 \pm 1.82$ | $6.36 \pm 1.83$ | 0.29 |
| SBRs putamen | $5.49 \pm 1.91$ | $4.71 \pm 1.99$ | 0.22 |

**Figure 1.**
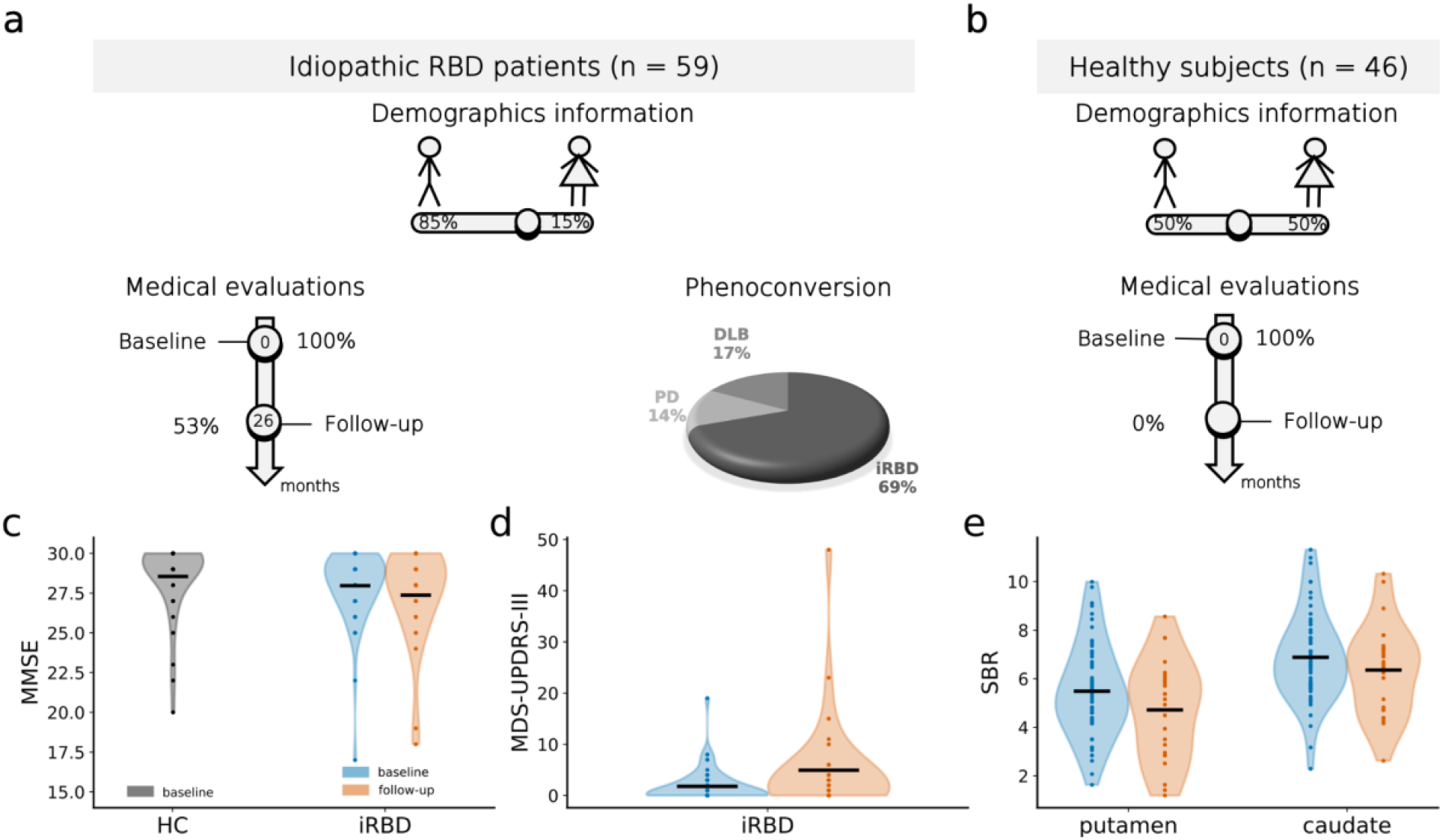
The main clinical and demographic data of idiopathic RBD (iRBD) patients - at baseline and follow-up - and HC subjects. Summary of sample characteristics of (a) iRBD patients and (b) age-matched healthy controls, including demographic information and percentage of subjects who underwent clinical and EEG assessment at baseline and first follow-up. The time-averaged (months) between baseline and follow-up is reported on the arrow. The pie graph represents the percentage of cRBD patients with PD (light gray) and DLB (grey), and stable iRBD (dark grey) for the whole dataset (n = 62). (c) Distribution of MMSE in iRBD patients at baseline (blue) and follow-up (orange) and in HC subjects (black). (d) Distribution of (d) MDS-UPDRS-III and (e) SBR (DaT levels) in the putamen and caudate in iRBD patients at baseline (blue) and follow-up (orange).

We excluded 3 people with iRBD and 2 healthy subjects from the following analysis due to excessive artifactual activity, which left less than 3 min of eye-closed resting-state data after cleaning. Moreover, 6 iRBD subjects did not undergo DaT-SPECT evaluation at baseline. For this reason, we excluded them when we investigated a possible correlation between EEG measures and DaT levels.

### Elevated LRTCs and bistability in neuronal oscillations characterize iRBD patients

To investigate whether a deviation of the operating point was already present in iRBD patients, we quantified the spectral profiles (30 Morlet wavelet decomposition) of (i) the LRTCs with DFA, (ii) the functional EI balance with fEI; and (iii) the neuronal bistability with BiS. Comparing the resting-state data of the two cohorts at baseline, we found that LRTCs and neuronal bistability were significantly (*p* < 0.05 - Wilcoxon rank sum test - BH correction) stronger in iRBD patients than in HC subjects in the delta-theta (2-7 Hz) band and beta-gamma (18-70 Hz) bands (Figure 2a,b). In contrast, iRBD patients showed a reduction (*p <* 0.05 *-* Wilcoxon rank sum test - BH correction) of fEI in the delta-theta band compared to HC subjects.

**Figure 2.**
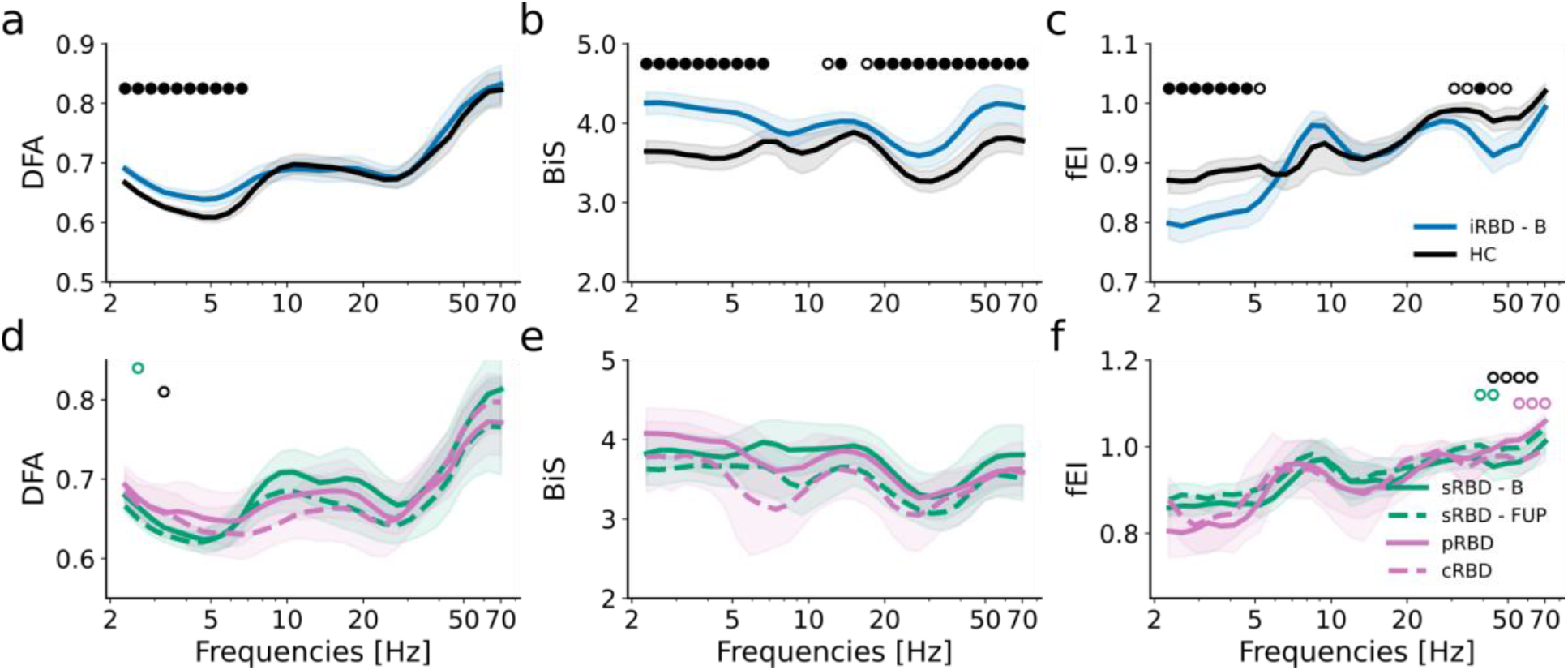
LRTCs and bistability are significantly stronger in iRBD patients at baseline than in healthy subjects. Group-level averaged of **(a)** DFA, **(b)** fEI, and **(c)** BiS for 59 iRBD patients at baseline (B - blue) and 46 healthy controls (black). Group-level averaged of **(d)** DFA, **(e)** fEI, and **(f)** BiS for 17 sRBD patients (green) and 11 cRBD patients (purple) at baseline (continuous line) and follow-up (FUP - dashed line). Shaded areas represent confidence intervals at 5% around the population mean (bootstrap, n = 1000). Empty circles highlight the frequency with a significant difference (p < 0.05) between populations using the Wilcoxon rank-sum test for pairwise differences: sRBD at baseline and follow-up (green circle); cRBD pre- and post-phenoconversion (purple circle); sRBD at follow-up and cRBD after phenoconversion (black circle). Filled circles highlight the frequency at which there is a statistical difference (p < 0.05) after the multiple comparison correction (Benjamini-Hochberg).

In the longitudinal analysis, people with iRBD exhibited no significant changes in these features from baseline to follow-up (Figure S1). No significant difference was found (Figure 2d-f) in sRBD between baseline and follow-up and in cRBD between pre- and post-phenoconversion after the BH correction. Moreover, we did not find any significant difference with phenoconversion after BH correction (Figure 2d-f).

These results suggested that iRBD affects the emerging brain dynamics mainly in the delta-theta band. The brain operating point is shifted toward the critical point (larger DFA) but still in an inhibition-dominated state (fEI < 1). Moreover, these metrics did not change over time, not even in patients developing the overt sign and symptoms of alpha-synucleinopathy. This result suggests that alterations of brain dynamics start early in the neurodegeneration process, but then they do not change further, possibly due to a floor effect.

### Low bistability and excitation-dominance are linked to nigro-striatal dopaminergic impairment

We first computed Spearman’s rank correlation coefficients between the putamen and caudate DaT levels (that is, the SBRs obtained from DaT-SPECT) and narrow-band neuronal fEI. The DaT levels were negatively correlated with fEI in the theta (5-7 Hz) band (Figure 3a). Moreover, cRBD patients (thick purple dots) exhibited higher values of theta band fEI (fEI > 0.87, f1-score: 52%, sensitivity: 72%, specificity: 54%) than the sRBD and pRBD (Figure 3b). DaT levels in the putamen were negatively correlated with fEI in the delta-theta band, mainly in the right centro-parietal electrode group (Figure 3d). In addition, we observed positive correlations, but not significant after BH correction, between EI (im)balance and motor dysfunctions (Figure 3c) in the theta (5-7 Hz) band and the gamma (> 30 Hz) band. We found no significant correlation between EI (im)balance and cognitive decline (Figure 3c).

**Figure 3.**
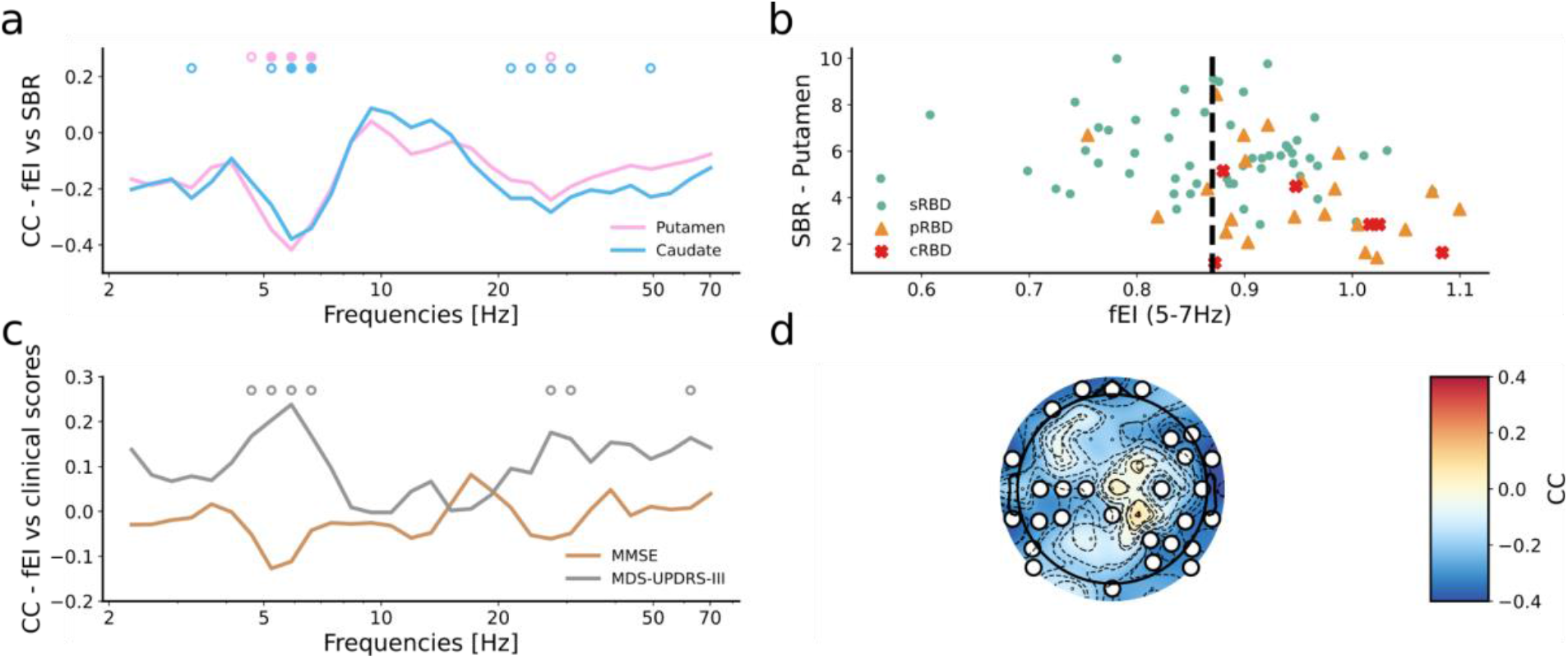
Functional EI ratio correlates with low DaT levels in the putamen. **(a)** Spearman’s correlation of averaged fEI across sensors with DaT levels in the putamen (pink) and caudate (azure). Empty circles highlight the frequency with a significant correlation (p < 0.05) between populations. The filled circles highlight the frequency at which there is a statistical correlation (p < 0.05) after the multiple comparison correction (Benjamini-Hochberg). **(b)** Scatter plot between DaT levels in the putamen and fEI in theta (5-7 Hz) band. Green circles show the stable iRBD patients (n=58), orange triangles show the progressive iRBD (n=n24), and red crosses show the cRBD patients (n=6). The dashed line represents the threshold value with the best phenoconversion prediction at baseline. **(c)** Spearman’s correlation of averaged fEI across sensors with MMSE (brown) and MDS-UPDRS-III (gray). Empty circles highlight the frequency with a significant correlation (p < 0.05) between populations. The filled circles highlight the frequency at which there is a statistical correlation (p < 0.05) after the multiple comparison correction (Benjamini-Hochberg). **(d)** Topoplot of the correlation coefficient between DaT levels and fEI in 5-7 Hz. White circles (or markers) show a significant correlation between the fEI computed for each electrode and DaT levels in the putamen after BH correction.

These results suggest a deviation of the operating point toward an excitation-dominated state in cRBD patients, which is the worsening of the biological disease severity (as expressed by the nigro-putaminal dopaminergic impairment) and tends to be associated with the clinical symptoms severity.

Subsequently, we investigated whether BiS is associated with DaT levels and clinical scores (Figure 4, S3). We found that DaT levels in putamen directly and significantly (*p* < 0.05, Spearman correlation coefficient - BH correction) correlated with bistability in the theta band (6-8 Hz), mainly in the central, parietal, and occipital electrode groups (Figure 4d). Moreover, we did not observe any correlation between BiS and MDS-UPDRS III or MMSE (Figure 4c). These findings suggest that iRBD patients with low bistability exhibit a significant nigro-striatal dopaminergic deafferentation, even if not paralleled by a clear association with clinical symptoms.

**Figure 4.**
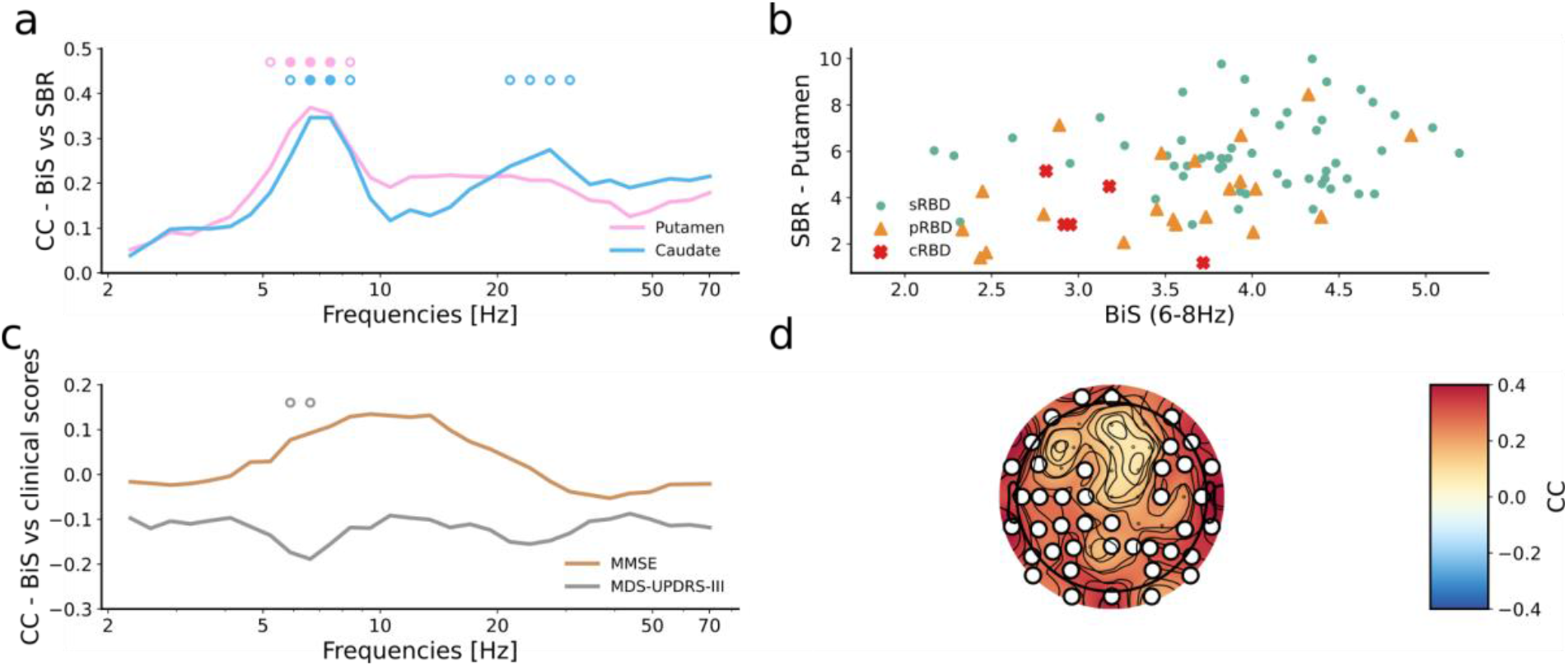
Low EEG bistability correlated with low DaT levels. **(a)** Spearman’s correlation of averaged BiS across sensors with DaT levels in the putamen (pink) and caudate (azure). Empty circles highlight the frequency with a significant correlation (p < 0.05) between populations. The filled circles highlight the frequency at which there is a statistical correlation (p < 0.05) after the multiple comparison correction (Benjamini-Hochberg). **(b)** Scatter plot between dopamine levels in the putamen and BiS in theta (6-8 Hz) band. Green circles show the stable iRBD patients (n=58), orange triangles show the progressive iRBD (n=24), and red crosses show the cRBD patients (n=6). The dashed line represents the threshold value with the best phenoconversion prediction at baseline. **(c)** Spearman’s correlation of averaged BiS across sensors with MMSE (brown) and MDS-UPDRS-III (gray). Empty circles highlight the frequency with a significant correlation (p < 0.05) between populations. The filled circles highlight the frequency at which there is a statistical correlation (p < 0.05) after the multiple comparison correction (Benjamini-Hochberg). **(d)** Topoplot of the correlation coefficient between DaT levels and BiS in 6-8 Hz. White circles (or markers) show a significant correlation between the BiS computed for each electrode and DaT levels in the putamen after BH correction.

Finally, we investigated the correlation of DFA values with DaT levels and clinical scores. DaT levels in putamen directly and significantly (*p* < 0.05, Spearman correlation coefficient - BH correction) correlated with the DFA exponent in the high-alpha (10-12 Hz) band (Figure 5a), mainly in the central, parietal, and occipital electrode groups (Figure 5d). In particular, cRBD patients exhibit low values of DFA exponent related to low DaT levels (Figure 5b), suggesting that low DFA exponent is related to biological disease severity. However, DFA and clinical scores only tended to be weakly correlated, without surviving correction for multiple comparisons (Figure 5c). Furthermore, there was no significant specific region-dependent correlation (Figure S4).

**Figure 5.**
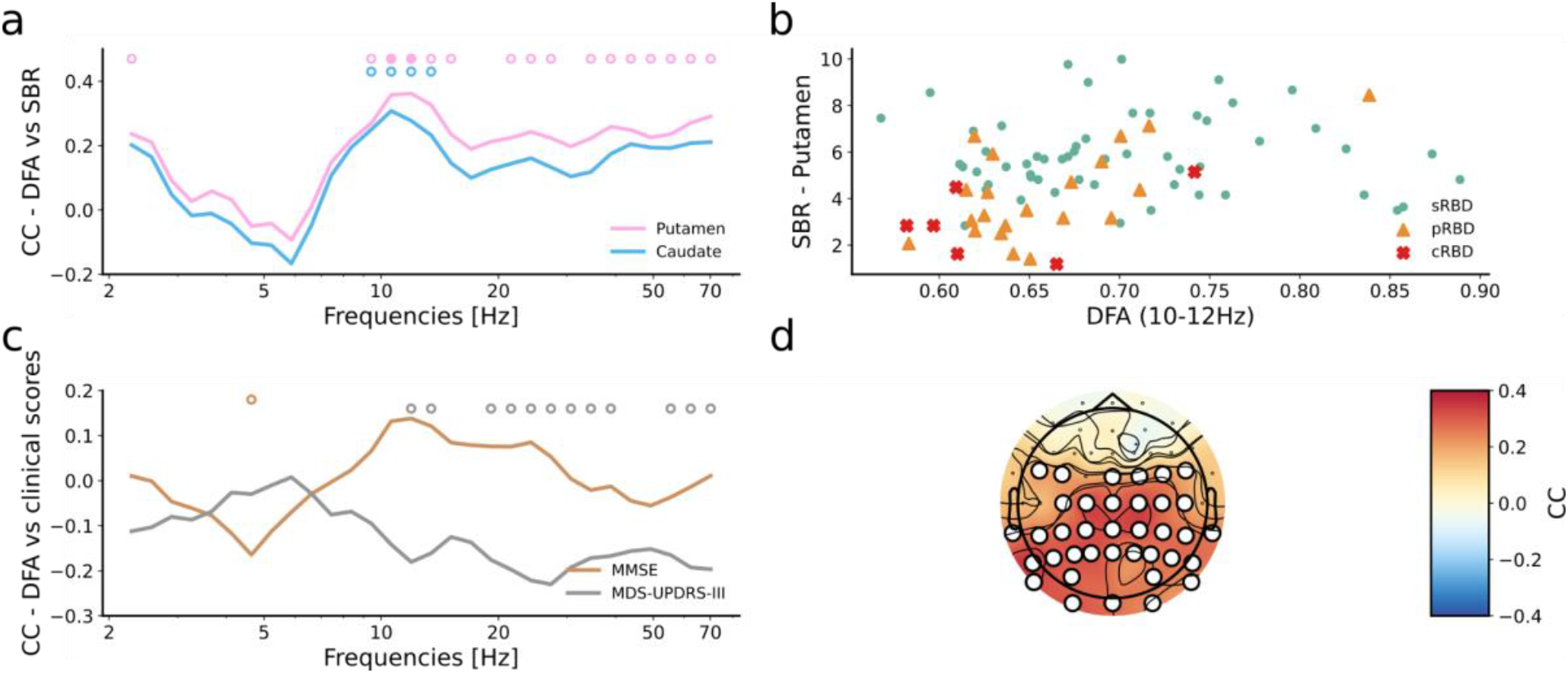
LRTCs positively correlate with nigro-striatal dopaminergic function. **(a)** Spearman’s correlation of averaged DFA across sensors with DaT levels in the putamen (pink) and caudate (azure). Empty circles highlight the frequency with a significant correlation (*p* < 0.05) between populations. The filled circles highlight the frequency at which there is a statistical correlation (*p* < 0.05) after the multiple comparison correction (Benjamini-Hochberg). **(b)** Scatter plot between DaT levels in the putamen and DFA in the alpha (10-12 Hz) band. Green circles show the stable iRBD patients (n=58), orange triangles show the progressive iRBD (n=24), and red crosses show the cRBD patients (n=6). The dashed line represents the threshold value with the best phenoconversion prediction at baseline. **(c)** Spearman’s correlation of averaged DFA across sensors with MMSE (brown) and MDS-UPDRS-III (gray). Empty circles highlight the frequency with a significant correlation (*p* < 0.05) between populations. The filled circles highlight the frequency at which there is a statistical correlation (*p* < 0.05) after the multiple comparison correction (Benjamini-Hochberg). **(d)** Topoplot of the correlation coefficient between DaT levels and DFA in 10-12 Hz. White circles (or markers) show a significant correlation between the DFA computed for each electrode and DaT levels in the putamen after BH correction.

### Phase synchronization positively correlates with the EI balance

The graph properties of synchrony networks were estimated using strength, clustering coefficient, and eigenvector centrality (see Method section).

EI balance and strength/clustering coefficient/eigenvector centrality were positively correlated (Spearman’s rank correlation, *p* < 0.05, BH corrected). In particular, the clustering coefficient was correlated with fEI in the theta-alpha (6-10 Hz) band (Figure 6a,b), and nodal strength with fEI in the theta-alpha (5-12 Hz) and low gamma bands (34-44 Hz) (Figure 6a,c,d). Nodal eigenvector centrality positively correlated with fEI at 18 Hz (Figure 6a). The strong correlation between phase synchronization and EI (imbalance), both hallmarks of brain disorders (Pusil 2019, Sunwoo 2017, Javed 2023), further suggest a transition toward a supercritical regime.

**Figure 6.**
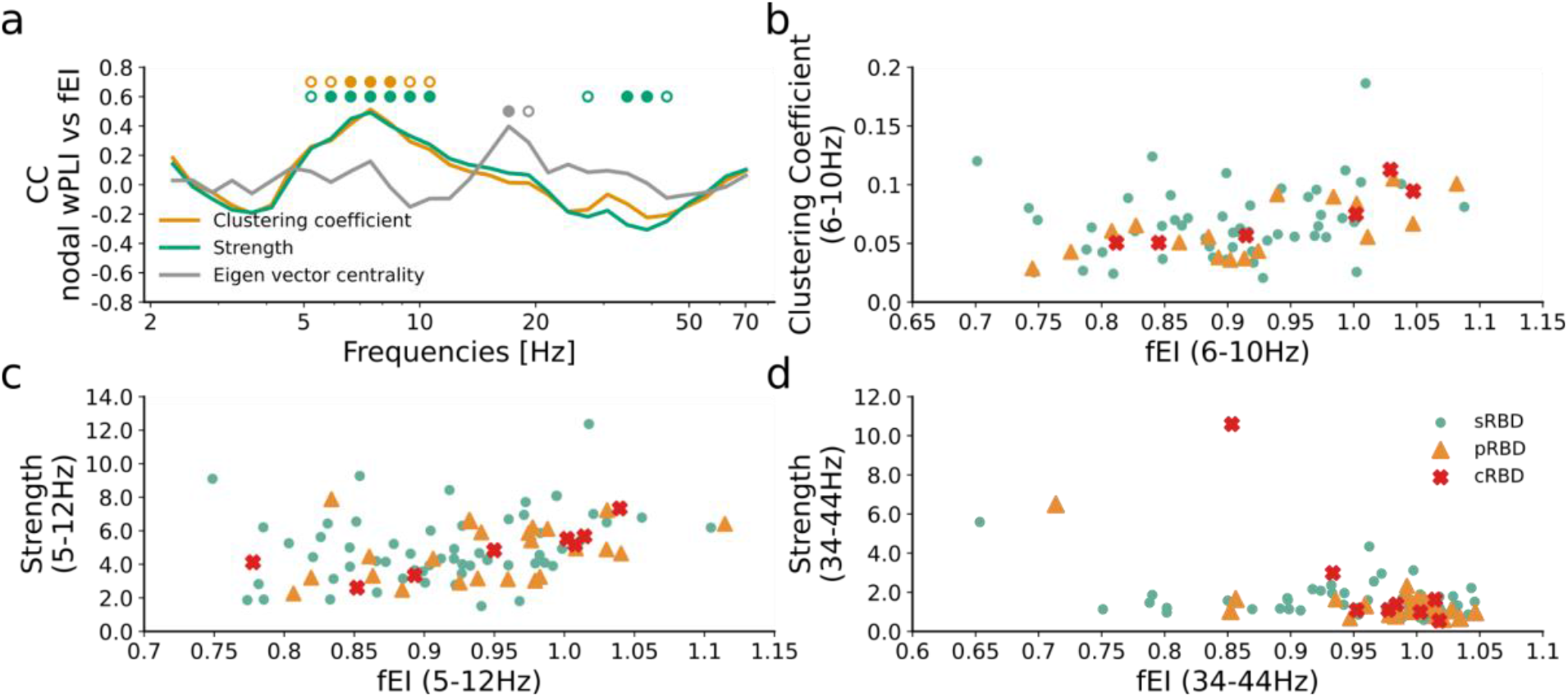
Phase synchronization correlates positively with EI balance. **(a)** Spearman’s correlation of fEI with nodal wPLI: clustering coefficient (yellow), strength (green), and eigenvector centrality (gray). Empty circles highlight the frequency with a significant difference (p < 0.05) between populations using the Wilcoxon rank-sum test for pairwise differences. The filled circles highlight the frequency at which there is a statistical difference (p < 0.05) after the multiple comparison correction (Benjamini-Hochberg). **(b)** Scatter plot of Clustering Coefficient and fEI in theta-alpha (6-10 Hz) band. Scatter plot of Strength and fEI in **(c)** theta-alpha band (5-12 Hz) and **(d)** gamma band (34-44 Hz). Green circles show the stable iRBD patients (n=58), orange triangles show the progressive iRBD (n = 24), and red crosses show the cRBD patients (n=6).

## Discussion

By longitudinally evaluating a group of patients with iRBD and healthy controls, we sought to understand how critical brain dynamics change throughout the prodromal phase of alpha-synucleinopathies to the overt stage of the disease. To this aim, we investigated 59 patients with iRBD and 46 age-matched HC subjects. We analyzed high-density EEG data by applying advanced quantitative analysis to extract several EEG features reflecting brain oscillations, under the conceptual framework of the critical brain hypothesis.

We found that spontaneous delta-theta band (5-7 Hz) activity of iRBD subjects exhibited higher bistability, higher long-range temporal correlations, and a smaller EI ratio compared to age-matched healthy controls. These findings suggest that iRBD localizes the brain operating point still in an inhibited state, with high levels of bistability.

In physiological conditions, the human brain works at the boundary of a critical phase transition between sub- and super-critical phases in a mild inhibited state (*i*.*e*., fEI close to 1) (Palva & Palva, 2018; Plenz, 2012; Priesemann et al., 2013; Wang et al., 2022). Recent modeling studies suggest that the critical point of the human brain occupies a wider range of the phase space, introducing the concept of the critical regime (Munoz, 2018; Wang et al., 2022) with a high degree of bistability (Wang et al., 2022). Here, we provided additional evidence to support the idea that bistable and critical dynamics are concurrent characteristics of oscillatory dynamics of the resting-state brain dynamics. Our results extended reports of altered critical dynamics of the human brain from known effects in epilepsy (Linkenkaer-Hansen et al., 2005), autism spectrum disorders (Bruining et al., 2020), and Alzheimer’s disease (Montez et al., 2009) since prodromal stages (Javed et al., 2022), also to prodromal stages of α-synucleinopathies.

As a second step, we evaluated the changes in brain oscillation metrics over time in a sub-group of enrolled iRBD patients with follow-up information, to gain insight into the longitudinal critical brain dynamics modifications in patients with alpha-synucleinopathies, since the prodromal phase. We found that iRBD showed a shift toward the critical point, by showing larger DFA, but still remaining in an inhibition-dominated state (*i*.*e*., fEI<1). Interestingly, these metrics did not change over time in our sample, not only in stable patients but also in those progressing to the overt stage of alpha-synucleinopathies. This result suggests that at least some of the critical brain dynamic modifications are an early event in the neurodegeneration cascade, but once altered, they do not change over time, possibly for a floor effect. Another possible explanation is that the limited number of phenoconverted patients reduced the statistical power of the analysis.

Subsequently, we investigated the association between the critical brain dynamics modifications and the biological and clinical severity of the disease. To this aim, we evaluated baseline and follow-up DaT-SPECT data, as a biomarker of biological disease severity, being correlated with DaT density. Moreover, we also analyzed longitudinal clinical modification of cognitive and motor symptoms. The EEG-derived criticality assessments were significantly correlated with nigro-striatal dopaminergic function, prominently in the bilateral-posterior channel groups. Despite the absence of significant modulation of critical brain dynamics in the longitudinal evaluation, the strong correlation between EI (im)balance and nigro-striatal dopaminergic impairment suggests that the critical brain dynamics of iRBD reflect the worsening of the biological disease severity. It is interesting to highlight that the EEG-derived metrics only tended to be associated with clinical disease severity, in particular with motor impairment, without surviving correction for multiple comparisons. This is not surprising. In fact, iRBD patients may have no to mild motor and cognitive symptoms for several years before phenoconversion (Joza et al., 2023). On the other hand, the nigro-striatal dopaminergic pathway may be altered several years before phenoconversion (Arnaldi et al., 2021), even in the absence of overt clinical symptoms (Iranzo et al., 2017). Thus, the biological damage caused by the degeneration may be already detectable in cortical (EEG-derived criticality measures) and subcortical (nigro-striatal dopaminergic function) structures, without a concomitant clear clinical impairment.

Notably, we observed that phenoconverted iRBD patients showed higher values of fEI and lower levels of DFA/BiS compared to stable iRBD patients at both baseline and follow-up visits. This evidence is further supported by a strong direct correlation between EI (im)balance and phase synchronization, both hallmarks for epilepsy (Fusca et al., 2022; Monto et al., 2007) and neurodegenerative diseases in geriatric subjects (Javed et al., 2022; Maestú et al., 2021; Montez et al., 2009; Pusil et al., 2019). Indeed, it is known that critical brain dynamics correlate with individual levels of phase synchronization (Fusca et al., 2022). As iRBD increases global phase synchronization (Roascio et al., 2022), we hypothesized that phase synchronization and EI balance were positively correlated in patients with iRBD. We estimated narrow band phase synchronization between EEG sensors using wPLI (Vinck et al., 2011), and these functional connectomes were treated as ‘graphs’ wherein each sensor was a ‘node’ cite (Bullmore & Sporns, 2009). In the presented data, we found a strong correlation between phase synchronization and EI (im)balance, both hallmarks of brain disorders (Javed et al., 2022; Pusil et al., 2019; Sunwoo et al., 2017), further suggesting a transition toward a supercritical regime. Therefore, even if no significant changes were demonstrated between baseline and follow-up assessments in our sample (perhaps due to the limited number of subjects), altogether these results suggest that once RBD patients phenoconvert to the overt stage of the disease (*i*.*e*., by expressing parkinsonism and/or dementia), they exhibit oscillatory dynamics of a system closer to the tip of the critical regime. Thus, the phenoconversion seems to change brain dynamics by further generating imbalance towards excitation-dominated conditions common to other brain disorders (O’Byrne & Jerbi, 2022; Zimmern, 2020).

We acknowledge that this work has some limitations. First, follow-up time was not fixed but ranged from 12 to 36 months approximately. Moreover, not all patients underwent instrumental (*i*.*e*., HDEEG and DaT-SPECT) follow-up. It has to be highlighted that the enrolled patients are currently still being followed. Thus, future studies on this sample would include longer follow-ups in all patients, hopefully with more time points. This will allow more accurate modeling of clinical and instrumental longitudinal changes. Moreover, this is a single-center study, with a limited number of subjects. Considering that longitudinal data from both clinical and advanced techniques are not easy to be collected, future multicenter studies with larger samples are needed to confirm the presented results. Finally, clinical longitudinal changes were assessed by using two metrics only (*i*.*e*., MMSE and MDS-UPDRS-III). Those two tools are commonly used as global measures of cognitive and motor function. However, it would be interesting to also evaluate the association between brain dynamics alteration and more comprehensive cognitive and motor function biomarkers, as well as with other signs and symptoms usually present in the alpha-synucleinopathies, such as autonomic function biomarkers.

In conclusion, this work shows for the first time that patients in the prodromal stage of alpha-synucleinopathies show cortical brain dynamic modifications by shifting the working point to excitation-dominated states getting closer to the overt stages of the disease. These results provide further evidence that both cortical and sub-cortical brain structures begin to be abnormal several years before the clinically overt syndrome (*i*.*e*., parkinsonism and/or dementia), thus confirming that patients with iRBD are in an incipient stage of alpha-synucleinopathy (Arnaldi & Mattioli, 2022). These results, on the one hand, increase our knowledge of the pathophysiological process underlying alpha-synucleinopathies since the prodromal stages. On the other hand, a better understanding of longitudinal cortical and sub-cortical modification along the continuum of alpha-synucleinopathies may provide new clues for innovative disease-modifying strategies.

## Acknowledgments

This work was carried out within the framework of the project “RAISE - Robotics and AI for Socio-economic Empowerment” and has been supported by European Union - NextGenerationEU. This work was developed within the framework of the DINOGMI Department of Excellence of MIUR 2018-2022 (legge 232 del 2016).

