## Supplementary materials for "Altered Brain Dynamics in idiopathic REM sleep behavior disorder: Implications for a continuum from prodromal to overt alpha-synucleinopathies"

#### $^{123}$I-FP-CIT-SPECT image acquisition and reconstruction.

To perform SPECT images, we used a dual-head Millennium VG camera (GE Healthcare) equipped with low energy, high resolution, parallel-beam collimators. Scans have been acquired 180-240 minutes after intravenous administration of 185 MBq of $^{123}$I-FP-CIT (DaTSCAN, GE Healthcare, Little Chalfont, Buckinghamshire, UK), and lasted 40 minutes. We applied a “step-and-shoot” protocol with a radius rotation lower than 15 cm, and 120 projections evenly spaced over 360° were generated. Total counts were comprised of between 2 and 3 million. We used an electronic zoom (zoom factor = 1.8) during the data-collecting phase to obtain an acquisition matrix’s pixel size of 2.4 mm. Subsequently, a digital zoom was applied during the reconstruction phase. The resulting images were sampled by cubic voxels (2.33 mm). We processed projections by applying an OSEM algorithm (8 interactions, 10 subsets) that included a pro back pair accounting for collimator blur and photon attenuation, and then post-filtering (3-D Gaussian filter with full-width at half maximum 58 mm). Photon attenuation was modeled with the approximation of a linear coefficient uniform inside the skull and equal to 0.11 $cm^{-1}$, and a 2D+ 1 approximation was applied in the simulation of the space simulation blur. Compensation for scatter was not performed.

#### Basal Ganglia software functioning.

Basal Ganglia software is an automatic algorithm based on a high-definition, 3-dimensional striatal template derived from the Talairach atlas. In detail, an automated algorithm performs fine adjustments in positioning blurred templates to match radioactive counts, meanwhile, it locates occipital regions of interest (ROI) to perform background evaluation. Also, a partial volume effect (PVE) correction is performed during the uptake computation of the putamen, caudate and background.

### Supplementary figures

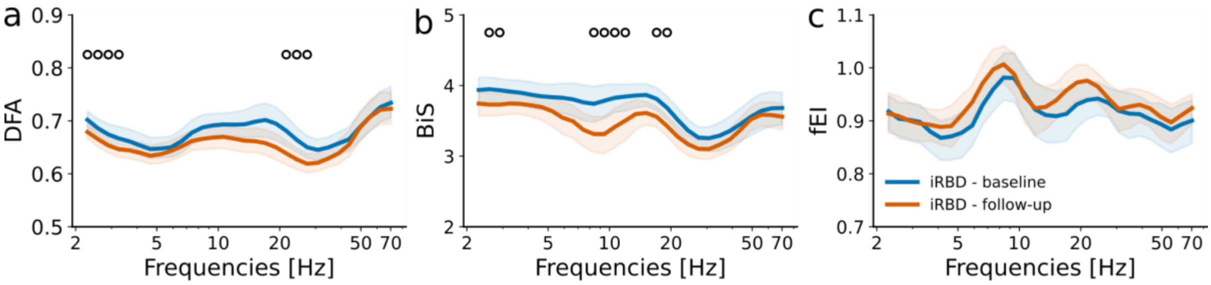


***Figure S1. LRTCs, EI balance, and bistability change with disease progression.*** Group-level averaged of DFA(a) and BiS (b) BiS for 31 iRBD patients at baseline (blue) and follow-up (orange); Shaded areas represent confidence intervals at 5% around the population mean (bootstrap, n = 1000). Black-empty circles highlight the frequency with a significant difference ($p < 0.05$) between populations using the Wilcoxon rank-sum test for pairwise differences. Black-filled circles highlight the frequency at which there is a statistical difference ($p < 0.05$) after the multiple comparison correction (Benjamini-Hochberg).
